# Strengthening interhemispheric connectivity increases interregional information transfer between human primary motor areas

**DOI:** 10.64898/2025.12.17.694630

**Authors:** Luca Tarasi, Arkadij Lobov, Cristina Del Prete, Domiziana Falaschi, Vito De Feo, Alejandra Sel

## Abstract

Communication between brain regions is thought to be supported by the temporal alignment of their oscillatory activity creating rhythmic windows of opportunity for efficient information transfer. We examined this possibility by manipulating the strength of coupling between the left primary motor cortex (lM1) and the right primary motor cortex (rM1) and examined the impact on activity transfer between the two areas measurable in the electroencephalogram (EEG) in 60 right-handed individuals. In three experiments we manipulated the strength of the pathway connecting the lM1 and rM1 by applying paired associative stimulation (ccPAS) using three different patterns, two of which increase interregional coupling strength. We measured the impact on EEG communication transfer in the alpha, beta, and theta bands at rest. Augmenting the strength of the lM1-to-rM1 pathway enhanced the interregional communication volume in the high-theta/low alpha band, and both volume and speed in the beta band, respectively. These changes, visible at rest, are related to changes in the transcallosal inhibitory influence of the lM1 over rM1 cortical excitability following ccPAS. By contrast, augmenting the rM1-to-lM1 route failed to modulate communication transfer in right-handed individuals. The results provide direct evidence of oscillatory intra-areal information transfer, demonstrating a link between motor network physiology and the resonant frequencies mediating its interactions.

## Introduction

Neural oscillations – rhythmic fluctuations of cortical activity – represent a fundamental mechanism underlying interregional communication at larger network scales (Buzsáki, 2006; Engel and Fries, 2010). In line with common accounts of neural communication (Fries, 2005, 2015), precise temporal alignment of the phase of such oscillatory rhythms is proposed to facilitate efficient information exchange across distant neural populations by enhancing transient windows of synaptic transmission. Recent evidence partially supports this notion showing that manipulating synaptic efficacy and connectivity in interregional connections with advanced protocols of brain stimulation can modulate oscillatory power (Sel et al., 2021) and phase synchronicity (Trajkovic et al., 2023) between two anatomically connected cortical regions. Despite these promising findings, it remains unclear whether enhanced interregional synchronous co-activations truly reflect information transfer in cortico-cortical communications. Here we directly tested this possibility in the human brain by using manipulations that either increase or decrease interregional synaptic plasticity between the left primary motor cortex (lM1) and the right primary motor cortex (rM1).

The pathway connecting the left and the right M1 is an ideal pathway in which to examine the effects of manipulating connection strength on interregional communication. It is well established that M1 represents a key structure supporting human voluntary movements. M1 is bilaterally represented and interconnected through dense bundles of callosal fibers that facilitate rapid interhemispheric exchange, dynamically regulating excitatory and inhibitory signals between the two homologous M1 regions, and ensuring smoothed coordination of bilateral actions (Hanajima et al., 2001; Meyer et al., 1998). The transcallosal pathway connecting lM1 and rM1 can be study in humans with transcranial magnetic stimulation (TMS): applying a conditioning TMS pulse over one M1 briefly suppresses motor-evoked potentials (MEPs) elicited from the test pulse in the contralateral M1, a phenomenon known as interhemispheric inhibition (IHI) (Ferbert et al., 1992). When this is done repetitively, the influence of one M1 over the other M1 is strengthened (Hernandez-Pavon et al., 2023; Rizzo et al., 2009, 2011). Such a procedure is often referred to as cortico-cortical paired associative stimulation (ccPAS), a non-invasive neuromodulation protocol based on spike-timing–dependent plasticity (STDP). ccPAS allows selective strengthening or weakening of cortical pathways by precisely controlling the temporal order of paired TMS pulses (Di Luzio et al., 2024). The impact of ccPAS on IHI can be visualized by examining MEP changes when paired TMS pulses are applied to the contralateral and ipsilateral M1. When this is done before and after ccPAS, a modulation of the IHI, and of the overall M1 excitability as measured by MEP changes, are observed (Hernandez-Pavon et al., 2023; Rizzo et al., 2009, 2011).

CcPAS may, therefore, be an ideal tool for investigating the impact of manipulating coupling between two cortical regions on the information transfer between them; if the effects of two different ccPAS protocols are compared then it should be possible to establish the impact of increasing connectivity in (1) the pathway connecting the lM1 with the rM1 (lM1-to-rM1 pathway) and (2) the reverse pathway connecting the rM1 with the lM1 (rM1-to-lM1 pathway), while holding constant the total amount of stimulation to each component area. In Experiment 1, we applied 15 minutes of ccPAS over lM1 followed by rM1 (lM1-to-rM1 ccPAS; each lM1 pulse was followed by a rM1 pulse) and measured changes in information transfer with electroencephalography (EEG). In Experiment 2, we investigated whether the changes in EEG information transfer were dependent on ccPAS stimulation order by reversing the order of ccPAS stimulation, i.e. applying the first paired pulse over rM1 and the second pulse over lM1, with the aim of manipulating coupling in the rM1-to-lM1 pathway. In both Experiment 1 and 2, the inter-pulse interval (lPI) between the two TMS pulses was set to 8ms corresponding to the optimal timing at which M1 exerts a physiological effect on the contralateral M1 by means of transcallosal connections (Ferbert et al., 1992; Rizzo et al., 2009). Lastly, in Experiment 3, TMS pulses were delivered over the lM1 and the rM1 almost simultaneously (1ms IPI – M1&M1 ccPAS); such a protocol does not induce pre- and post-synaptic activation and thus is not expected to induce Hebbian-like changes in the M1-M1 interhemispheric connections acting as control condition. Exactly the same number of pulses were applied to LM1 and RM1 in all 3 Experiments (Figure 1), carefully controlling for the impact on each component area when altering the strength of the pathway between them.

**Figure 1:**
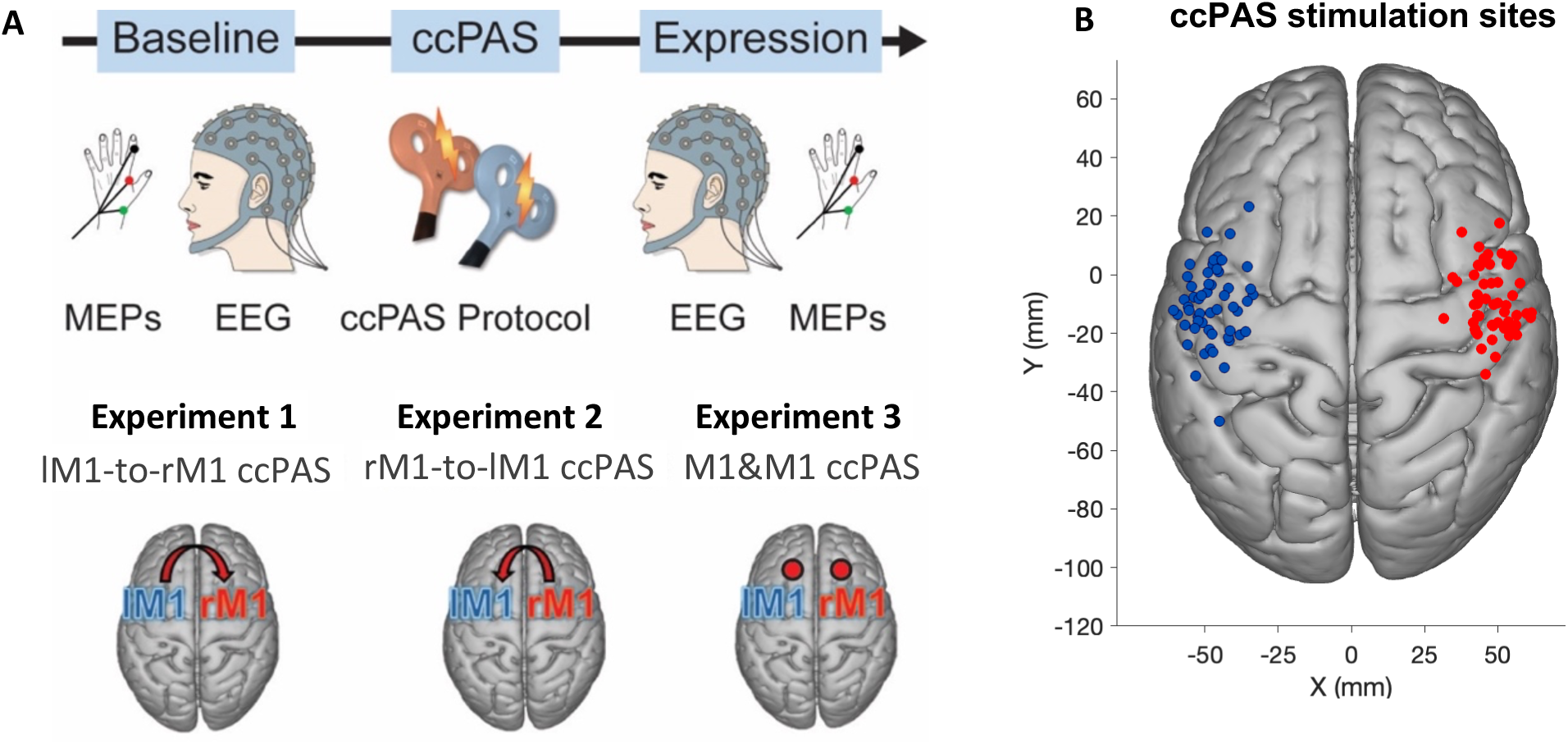
**A.** Experimental design for Groups 1, 2 and 3. **B.** Individual scalp hotspots for the right primary motor cortex (rM1) and left primary motor cortex (lM1). The filled circles in red (rM1) and blue (lM1) represent the subject scalp hotspot for Group 1, 2 and 3 in standardized MNI space

Before and after ccPAS, we recorded resting state EEG activity from fronto-central and centro-parietal areas where previous beta, alpha and theta oscillatory responses linked to motor interhemispheric communication are observed (Barone and Rossiter, 2021; Donner et al., 2009; Engel and Fries, 2010; Pscherer et al., 2019, 2023; Sauseng et al., 2009; Tempel et al., 2020). To establish the impact of ccPAS on oscillatory activity, we first measured changes in phase coherence between the left and right M1 using the weighted phase-lag index (wPLI), an index that quantifies EEG synchrony while minimizing confounds due to volume conduction (Vinck et al., 2011). Crucially, to assess changes in the directionality of information flow, we employed cross-correlation analysis. These allowed us to measure alterations in the amplitude and latency of inter-areal transmission, enabling separate evaluation of its strength and direction (Adhikari et al., 2010). Concurrently, we examined changes in interhemispheric inhibition (IHI) and overall M1 cortical excitability by recording MEPs to single or paired TMS pulses over left and right M1 (Figure 1).

The examination of directionality changes in resting EEG activity enabled us to look at a range of oscillatory effects in the EEG. Different frequency bands have been consistently linked to distinct functional roles within the human motor system; thus, beta-band rhythms (14–30 Hz) are typically associated with interhemispheric synchronization during bimanual and unimanual tasks (Murthy and Fetz, 1996), as well as to motor execution, motor preparation and the stable maintenance of motor states (Barone and Rossiter, 2021; Donner et al., 2009; Engel and Fries, 2010; Tarasi et al., 2025). Alpha oscillations (8-13) represent the dominant oscillatory activation in the sensorimotor cortex in the absence of movement (Haegens et al., 2011; Stefanou et al., 2018), and movement initiation and cessation are linked to increases and decreases in theta band (4-7Hz), respectively (Harper et al., 2014; Picazio et al., 2014; Tsujimoto et al., 2010; Zhang et al., 2008).

## Results

In two experiments with a total of 40 right-handed individuals (20 per experiment), we investigated, respectively, whether increasing coupling in the route connecting the left M1 to the right M1, or the pathway from the right M1 to the left M1 led to modulation of interareal information transmission between the two primary motor cortices. We contrasted the effects of the two types of ccPAS, repeated paired stimulation of lM1 followed by rM1 (Experiment 1) or, vice versa, rM1 followed by lM1 (Experiment 2) on time-frequency EEG oscillatory responses, recoded during rest. We focused on motor-relevant frequency bands theta, alpha and beta (4-30Hz) in fronto-central and centro-partietal electrodes bilaterally (Methods: EEG recording and analysis) known to reflect interhemispheric communications between human motor cortices during rest (Stefanou et al., 2018). In addition, we tested for non-specific effects of the ccPAS in Experiment 3 (20 right-handed individuals), where the pulses involved in ccPAS were delivered almost simultaneously (1ms IPI) (Figure 1).

Prior to examining the changes in communication information transfer led by ccPAS, we performed a series of analysis to investigate changes in M1 cortical excitability focusing on the inhibitory influence of the contralateral M1 over the ipsilateral M1 – i.e. interhemispheric inhibition (IHI) (Hernandez-Pavon et al., 2023; Rizzo et al., 2009, 2011) (Method section: MEP analysis). First, we compared the MEP amplitude changes when we applied either a single pulse TMS (spTMS) over the ipsilateral M1 or a paired-pulse TMS (ppTMS) over the contralateral M1 (conditioning pulse) followed by the ipsilateral M1 (test pulse). We recorded MEPs from the left first dorsal interosseus (FDI) muscle at rest. We demonstrated that the conditioning pulse delivered in the contralateral M1 did indeed reduce the impact of the ipsilateral M1 pulses in comparison to the impact of spTMS on the same component area; this occurred in both the left and the right M1 confirming the long-standing phenomenon of IHI (Rizzo et al., 2009, 2011) (Figure S1). Second, we examined the impact of repeatedly inducing lM1 activity just before inducing activity in rM1, or vice versa, during ccPAS. Again, we did this by measuring MEPs recorded in response to spTMS over M1 either alone or preceded by a conditioning TMS pulse in the contralateral M1, but we did so before and after a 15-minute period of ccPAS. Here we demonstrated that we could manipulate the connectivity in the transcallosal pathways between the two primary motor cortices. The three ccPAS protocols used in the three Experiments exerted distinct effects on interhemispheric inhibition of M1 activity (F2,57 = 3.24, p = .046, n2p = .0.1). Specifically, the lM1-to-rM1 ccPAS significantly increased the inhibitory effect of the contralateral hemisphere on ipsilateral M1 excitability – i.e. potentiation of IHI (F1,19 = 6.47, p = .020, n2p = 0.25); these changes did not appear immediately after ccPAS (i.e. during the first 20 trials after the ccPAS protocol – p = 0.32), but evolved with time emerging around 20 minutes after the stimulation during the last 20 trials of the block (t1,19 = 2.65, p = 0.016, d = 0.34), in line with previous investigations (Chiappini et al., 2022; Rizzo et al., 2009, 2011) (Method section: MEP analysis). By contrast, the rM1-to-lM1 led to a decreased IHI effect on the ipsilateral M1, this is greater M1 excitability when the test pulse was preceded by a conditioning pulse on the contralateral M1 (F1,19 = 8.02, p = .011, n2p = .0.3) (Figure S1) (Rizzo et al., 2009, 2011). Lastly, in Experiment 3 where the ccPAS were delivered almost simultaneously in a manner that does not follow Hebbian principles, we do not observe any changes in IHI as measured by M1 MEP amplitude changes (p = 0.75).

After determining the impact of ccPAS in the connectivity strength between the pathways interconnecting the left and right M1, we then assessed the changes on their inter-site communication information transfer. We started by examining whether manipulating the connectivity strength of the M1-M1 callosal tracts changes their inter-areal phase coupling. Subsequently, we examined whether the changes in phase coupling synchronization are accompanied by changes in the transmission of information between the two nodes of the examined network at rest. Specifically, to assess changes in intra-areal coherence we used the weighted phase lag index (wPLI): a phase lag-based measure not affected by volume conduction and not biased by the sample size (Tarasi et al., 2022; Trajkovic et al., 2023). We examined a cluster of electrodes located around the areas of stimulation in the right and left hemisphere spanning homologous M1 regions, including frontal, fronto-central, central and centro-parietal sensors. The wPLI was evaluated for all channel pairs within the ROIs in both the right and left hemisphere with the intention to explore interhemispheric connections (Figure 2 & 3). Subsequently, non-parametric cluster-based permutation analysis was used to compare wPLI before and after ccPAS stimulation between different pairs of sensors of ROIs in distinct frequency bands (for details, see Methods).

**Figure 2:**
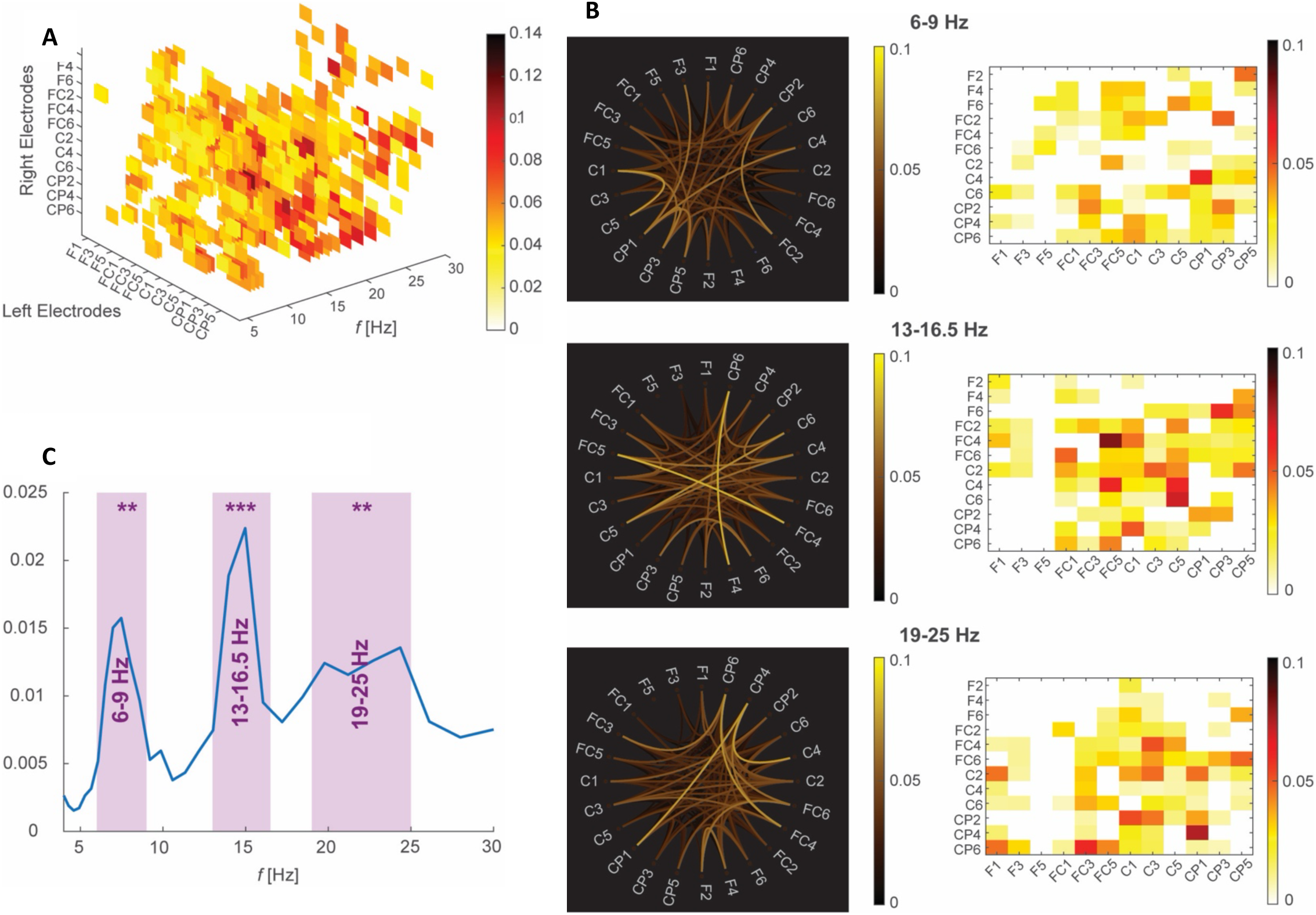
Effects of left M1-to-right M1 ccPAS over interregional connectivity Changes in wPLI Before vs After ccPAS. Changes in interregional coupling between the lM1 and the rM1 (N=17). **A.** 3-D representation of the connectivity strength changes measured by wPLI in the sensors of interest, and across all frequency bands in the group undergoing lM1-to-rM1 ccPAS. Each colored square represents the normalized wPLI difference when contrasting the activity recorded in the Expression block vs activity recorded in the Baseline block for each pair of electrodes and for each frequency. The normalization was performed by dividing the wPLI difference between activity recorded in the Expression vs the Baseline by the value of the wPLI recorded at Baseline. Only significant clusters are shown. **B.** Representation of the wPLI changes across all relevant electrodes in the selected frequencies of interest for theta, low alpha and low and high beta. **C.** Averaged wPLI values across all the electrode pairs showing a significant increase when comparing the Baseline and Expression blocks. Shaded areas in purple indicate the frequency bands showing greatest wPLI changes selected for further cross-correlation analysis. ** denotes significant results with a p value <0.01

**Figure 3:**
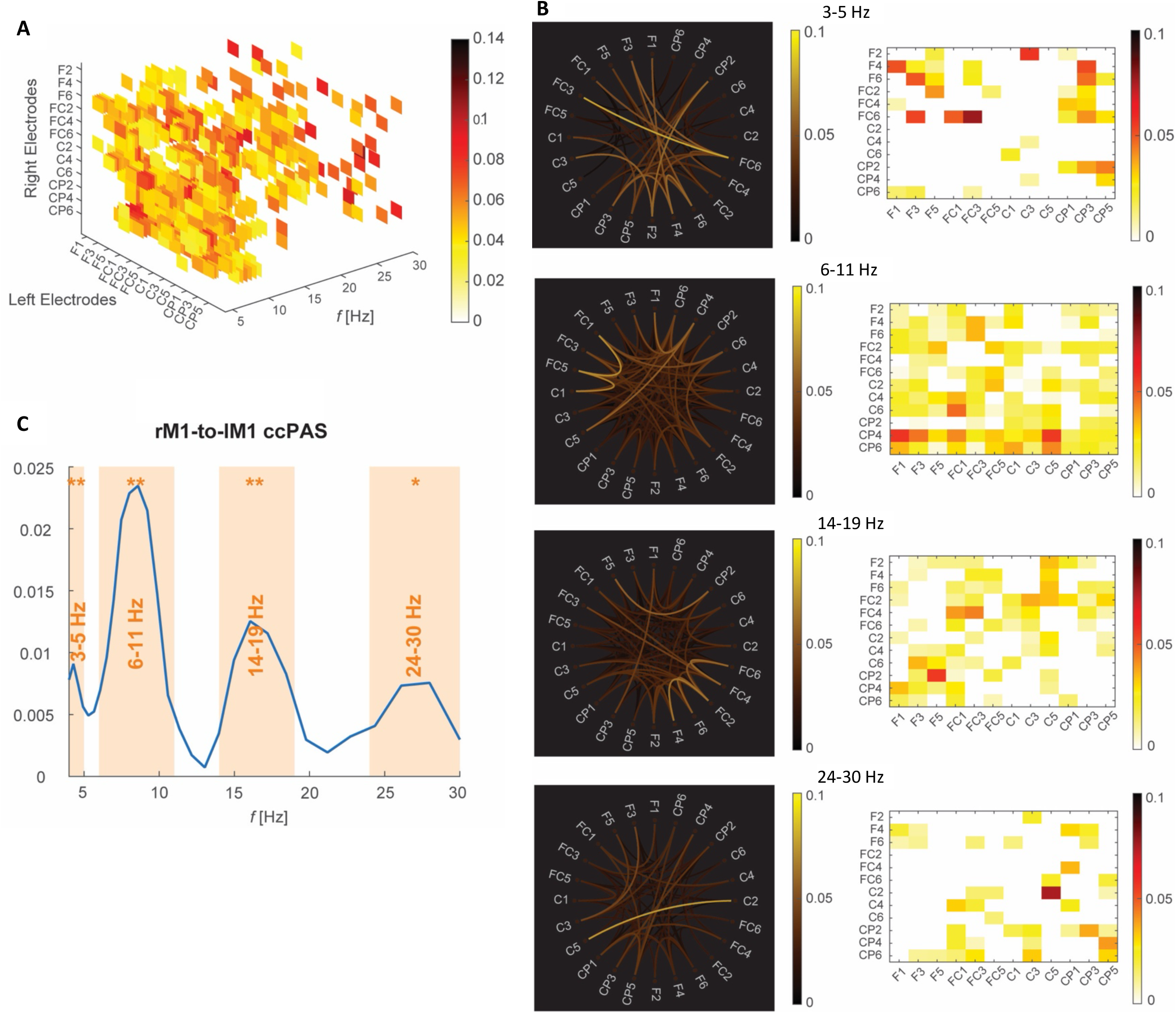
Effects of right M1-to-left M1 ccPAS over interregional connectivity Changes in wPLI Before vs After ccPAS. Changes in interregional coupling between the rM1 and the lM1 (N=18). **A.** 3-D representation of the connectivity strength changes measured by wPLI in the sensors of interest, and across all frequency bands in the group undergoing rM1-to-lM1 ccPAS. Each colored square represents the normalized wPLI difference when contrasting the activity recorded in the Expression block vs activity recorded in the Baseline block for each pair of electrodes and for each frequency. The normalization was performed by dividing the wPLI difference between activity recorded in the Expression vs the Baseline by the value of the wPLI recorded at Baseline. Only significant clusters are shown. **B.** Representation of the wPLI changes across all relevant electrodes in the selected frequencies of interest for theta, low alpha and low and high beta. **C.** Averaged wPLI values across all the electrode pairs showing a significant increase when comparing the Baseline and Expression blocks. Shaded areas in orange indicate the frequency bands showing greatest wPLI changes selected for further cross-correlation analysis. * and ** denotes significant results with a p value <0.05 and <0.01, respectively

The results of the wPLI analysis indicate that the frequency and synchronicity of connectivity changes are dissociable for the two ccPAS protocols delivered in Experiment 1 (Figure 2) and Experiment 2 (Figure 3). Specifically, in Experiment 1 (first pulse delivered to the left M1 and followed by a pulse on the right M1), there was an increase in the connectivity in the slow frequencies, i.e. the cross-frequency range high theta / low alpha (i.e. 6-9Hz), and in the faster frequencies: low (13-16.5Hz) and high (19-25Hz) beta frequency range (p = 0.0120). On the other hand, in Experiment 2 (rM1-to-lM1 ccPAS) the connectivity changes were circumscribed to the following frequency ranges: 3-5Hz, 6-11Hz, 14-19Hz and 24-30Hz (p = 0.0172). By contrast, in Experiment 3 (where both TMS pulses were delivered almost simultaneously, i.e. 1ms inter-pulse interval) the analysis did not show any differences in wPLI between the Baseline and Expression blocks, indicating a lack of connectivity changes in the M1-M1 ccPAS participant group. These changes in coherence in Experiment 1 and 2 were not accompanied by changes in the amplitude (power) of the signal in any of the frequency bands (all p values > 0.05).

Next, we proceeded to directly measure any changes in inter-site communication information transfer between the right and the left M1. We focused on the frequency bands where the greatest interregional coupling changes were found for each experiment. To select these frequency ranges, we started by computing the average difference of wPLI values (calculated by subtracting the Baseline wPLI values from the expression wPLI values) across all the electrode pairs. For those electrode pairs in which the wPLI values did not significantly differ between the Baseline and the Expression blocks, the average difference value was set to zero. This was done for each frequency point within the frequency bands where a significant increase in synchronicity as measured by wPLI changes was identified (Sup. Figure 2). This procedure was done separately for the Experiment 1 and the Experiment 2 (see the Methods section for a detailed description). Specifically, in Experiment 1 a Friedman test showed a significant difference between the selected frequency bands and the remaining frequency bands (χ²(3, N = 17) = 25.729, p <0.001). Post hoc Wilcoxon signed-rank tests with False Discovery Rate correction indicated that all selected frequency bands differed significantly from baseline (6-9Hz: Z=3.101, p=0.002; 13-16.5Hz: Z=3.621, p=0.000; 19-25Hz: Z=3.479, p=0.0015). Similarly, the Friedman test performed in Experiment 2, showed a significant difference between the selected frequency bands and the remaining frequency bands (χ²(4, N = 14) = 21.543, *p* <0.001). Post hoc Wilcoxon signed-rank tests with False Discovery Rate correction indicated that three of the four frequency bands analysed differed significantly from baseline (3-5Hz: Z=2.678, p=0.0093; 6-11Hz: Z=3.027, p=0.004; 14-19Hz: Z=3.593, p=0.004; 24-30Hz: Z=2.199, p=0.028). These frequency bands were the greatest change in synchronicity – i.e. changes in averaged wPLI values across sensors, between before and after the ccPAS intervention are observed were taken for further analysis.

For each identified frequency range, we then computed the cross-correlation between each of the pairs of sensors in the ROI for each participant separately in both experimental groups. First, we measured the correlation between the envelope of two signals recorded in each electrode pair with the aim to examine the directionality of the information flow. Specifically, we extracted three activity peaks in the signal: a central peak indexing the volume conductance, a left peak representing changes in the information transfer from sensors located under or adjacent to the rM1 to their homologous electrode pair located near the lM1, and a right peak indexing information transfer from sensors near the lM1 to their corresponding electrode under or adjacent to the rM1 (Figure 4A). The analysis focused on the differences in peak amplitude and latency (lag) between the Baseline and the Experimental blocks, for each of the experimental groups independently. In the participant group where the lM1 pulse preceded the rM1 pulse (lM1-to-rM1 ccPAS, i.e. Experiment 1) we observe a significant increase in the right peak amplitude in several electrode pairs indicating an increased volume of information transfer from the left to the right M1 cortex in the Expression vs the Baseline block. This information transfer occurred across all the analyzed frequency windows: 6-9Hz (p: 0.0324), 13-16.5Hz (p: 0.0224) and 19-25Hz (p: 0.0204), with a particularly enhanced volume of information transfer in the slower frequencies (6-9Hz) (Figure 4A). Moreover, the results show a decrease in the lag of the right peak in a number of electrode pairs circumscribed to the higher beta frequencies 19-25Hz (p: 0.028), indexing faster information transfer in the Expression vs the Baseline blocks. Additionally, there is a smaller increase in the information transfer from the sensors located around the right M1 to the sensors around the left M1, as indicated by a significant left peak amplitude increase in the 6-9Hz frequency window (p: 0.0444) (Figure 4A). By contrast, the cross-correlation analysis performed in the participant group that underwent the rM1-to-lM1 ccPAS – i.e. Experiment 2, did not show any significant changes in information transfer when contrasting the amplitude and lag of the right and the left peaks recorded in the Expression vs the Baseline blocks in the selected frequency windows (p > 0.05).

**Figure 4:**
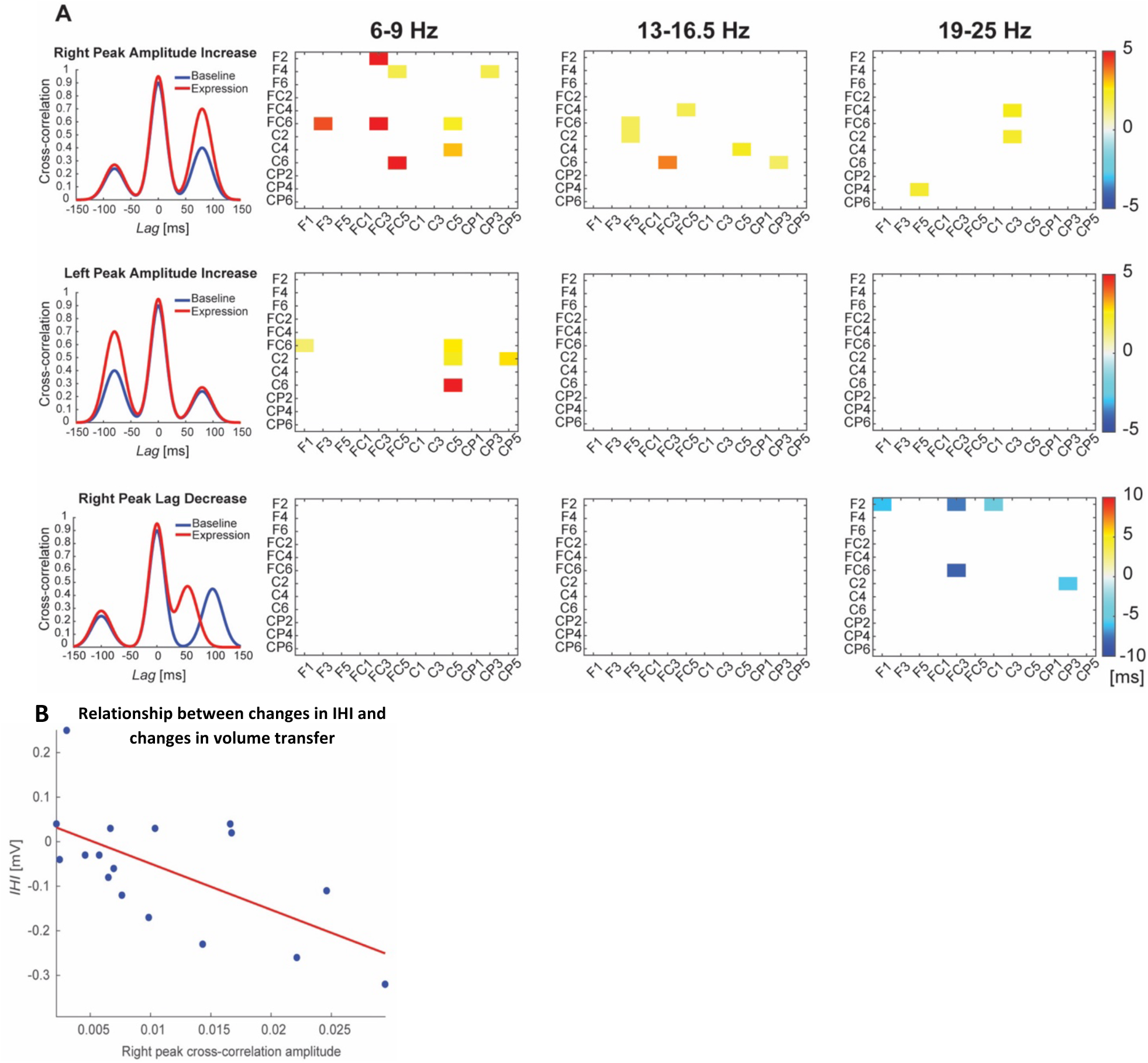
Effects of ccPAS over the left M1 and right M1 on neural activity transfer Changes in cross-correlation amplitude and latency Before vs After lM1-to-rM1 ccPAS. **A.** Representation of the significant increase in peak amplitude (top and middle row) and decrease peak latency (bottom row) in cross-correlation measures when comparting the Baseline and the Expression blocks for each electrode pair in the participant group undergoing lM1-to-rM1 ccPAS (N=17). Warm colors indicate an increase of cross-correlation values in the Expression block vs the Baseline block, cold colors indicate a decrease of cross-correlation values between Expression vs Baseline; the white squares indicate non-significant differences. **B.** Pearsons correlation between the changes in information volume between electrodes C4 and C5 in the 6-9Hz frequency range, and changes in the interhemispheric inhibition in the participant group where the ccPAS protocol target the pathway connecting the lM1 with the rM1.

Furthermore, we explored whether the increases in the volume and speed of information transfer found in Experiment 1 (participants undergoing lM1-to-rM1 ccPAS) were associated with the increases in interhemispheric inhibition (IHI) observed in that same group when comparing the Expression vs Baseline blocks. For these analyses we selected the electrode pair showing the greatest enhancement in information volume transfer and information speed transfer for each of the frequency windows analyzed (i.e. C2-C3 and C4-C5) and computed the difference value of peak amplitude and peak latency changes (Expression block – Baseline block). Similarly, we took the difference IHI value computed as the difference between the IHI recorded in the Expression block minus the IHI recorded in the Baseline block. A Pearson correlation of these difference values showed that the greater the information volume transfer between the C4 and C5 electrodes (sensors directly located over homologous M1 areas) in the 6-9Hz frequency window, the stronger the increase of IHI in the participant group undergoing lM1-to-rM1 ccPAS (r = -0.6127, p=0.0267) (Figure 4B). No other significant correlations were found within the values explored (p > 0.05).

## Discussion

The synchronization of intra-areal neuronal oscillations is generally regarded as a key element in communication between brain regions, with information transfer being channeled via distinct frequency bands (Fries, 2005, 2015; Salinas and Sejnowski, 2001). Converging theories suggest that interregional information transfer occurs when oscillations in different neuronal sets create rhythmic opportunities allowing for the synchronous orchestration of action potentials across those neuronal cells (Fries, 2005, 2015; Salinas and Sejnowski, 2001). It has also been posited that the efficiency (volume and speed) of communication information relies on the synaptic efficacy of connections between such neuronal groups (Fries, 2015; Horrocks et al., 2024; van Ede et al., 2018). Here we directly tested this hypothesis in the human brain by using manipulations that have been established to modify connectivity strength in a human cortico-cortical pathway, the route connecting the left and the right M1 (Hernandez-Pavon, 2025; Rizzo et al., 2009, 2011). We demonstrate that changing short-term synaptic efficacy of the pathway connecting the left M1 with the right M1 changes the volume and speed of information between these two areas.

The repeated application of TMS pulses to the left M1 followed by the right M1, lM1-to-rM1 ccPAS, evokes synchronous pre- and post-synaptic activity in the M1-M1 pathway. This results in increases of the volume and the speed of communication information between the two targeted nodes, that may also extend towards interconnected frontal and parietal regions. Specifically, the information transfer from the left to the right M1 cortex in neural activity recorded over frontal, central and centro-parietal sensors adjacent to M1 cortices improved following lM1-to-rM1 ccPAS. These changes are present across a wide range of oscillatory bands including high theta, low alpha and beta ranges, but they are particularly strong in the high theta band. Similarly, the speed of information transfer from the lM1 to the rM1 also increased after strengthening the route connecting these two regions, particularly in the high beta frequencies. Improvements in information transfer have been linked to stronger or more efficient synaptic connections in rodents (Hartung et al., 2016a, 2016b; Siegle et al., 2021). In this vein, targeting the lM1-rM1 route with ccPAS may lead to enhanced functional efficiency. This could occur through mechanisms such as an increase in the number of receptors on a synaptic membrane, a process consistent with the principles of long-term potentiation (LTP) (Abraham et al., 2024). This is consistent with previous evidence demonstrating ccPAS-induced changes in functional connectivity in the motor control network (Johnen et al., 2015; Lazari et al., 2022). Notably, we also observe increases in quantity of information in the reversed right M1 to left M1 direction, specifically in the high beta band. These changes could be related to a rebound of information from the right M1 to the left M1 through the bidirectional M1-M1 pathway, in response to the marked changes in information volume observed from the left to the right M1 cortex in the high beta band (Watanabe et al., 2014). Overall, we demonstrate that increasing the short-term synaptic efficacy of the lM1-rM1 pathway changes the volume and speed of information transfer between the left and the right M1 regions in a group of right-handed individuals.

Interhemispheric inhibition in the motor system is essential for coordinating and controlling voluntary movements. These reciprocal interactions are subserved by interhemispheric synchronization of oscillatory neuronal activity that resonate at different frequencies (Stefanou et al., 2018). Different cortical rhythms in the beta, alpha, and theta ranges are associated with distinct functional roles in motor control; these oscillatory response patterns are instrumental for the dynamic interplay between the two primary motor cortices supporting smooth, coordinated motor performance (Picazio et al., 2014; Stefanou et al., 2018). Beta oscillatory activity is a dominant frequency band in interregional communication in the motor control circuit, particularly during inhibition and absence of movement (Ferreri et al., 2014; Picazio et al., 2014; Zhang et al., 2008). In non-human primates, 20–40 Hz oscillations in the primary motor cortices (M1) of both hemispheres transiently synchronize during bimanual and unimanual motor tasks; in humans, low-beta band interhemispheric communication plays a critical role in regulating movement, and this process is notably influenced by hand dominance (Serrien and Spapé, 2009). Furthermore, theta (4-7 Hz) associated with movement initiation and cessation (Harper et al., 2014; Picazio et al., 2014; Tsujimoto et al., 2010; Zhang et al., 2008). By contrast, alpha oscillatory activity represents the dominant rhythm at rest in the sensorimotor cortex (Haegens et al., 2011). Here we observe that manipulating the strength of connectivity in the route connecting the left M1 with the right M1 increases the volume of information transfer in the cross-frequency theta/alpha range (6-9Hz), the low beta range (13-16.5Hz) and the high beta range (19-25Hz). These results indicate that the neuronal architecture supporting lM1-to-rM1 interhemispheric communication has fundamental resonant properties in different frequency bands and that even in the absence of any motor task, different manipulations (left-to-right vs right-to-left direction) of the M1-M1 pathway strength affect specific communication channels. It is worth noting that the functional impact of changes in the different channels may putatively be most apparent during different aspects of motor behavior, such as unimanual and bimanual coordination tasks.

By contrast, reversing the order of stimulation so that right M1 pulses are followed by left M1 pulses during the repeated stimulation protocols is not accompanied by changes in information transfer. While rM1-to-lM1 leads to an increase in oscillatory synchronicity across a range of frequencies including theta proper (3-5Hz), cross-frequency high theta / alpha (6-11 Hz), low beta (15-18Hz) and high beta (26-32Hz), no changes in amount or speed of information transfer are observed. It is well-established that the motor system organization varies with handedness, and that the callosal size can vary with the degree of handedness. For example, less strongly handed individuals have a larger callosa in both the posterior and the anterior midbody regions (Luders et al., 2010). In the current study, we tested right-handed participants with a strong handedness, which means that there is likely to be an asymmetry between the routes connecting the right M1 and the left M1. Accordingly, the manipulation of these two pathways with ccPAS lead to divergent results, whereby only strengthening the lM1-to-rM1 pathway increases the volume and speed of information transfer, making this route more efficient.

These results are in line with previous evidence suggesting that the degree of laterality influences the transcallosal interhemispheric transfer speed as measured by the Poffenberger paradigm (Bernard et al., 2011). They also accord with similar directional effects found when manipulating cortico-cortical pathways in other brain networks like the visual cortices, where it is shown that reinforcing the left V5-to-right V5 pathway (vs the reversed stimulation direction) heightened sensitivity to horizontal visual motion (Chiappini et al., 2022). Notably, while the rM1-to-lM1 ccPAS protocol strengthens the connections from the right to the left M1, it also stimulates the lM1-to-rM1 route in a way that activates the post-synaptic neurons before the pre-synaptic neurons. According to the principles of Hebbian plasticity, this activation timing leads to long-term depression (LTD), making such connections weaker. Therefore, the asymmetrical pattern of results observed in Experiment 1 and Experiment 2 may be explained by concurrent influence of LTP and LTD mechanisms, which express themselves differently due to the inherent asymmetry of the two targeted cortical pathways. We demonstrate that the nature of the interregional pathways directly influences the volume and speed of information transfer, revealing a causal link between physiological asymmetries and oscillatory synchronization within the motor control system.

Furthermore, we analyzed the influence of manipulating the M1-M1 cortical pathways on M1 cortical excitability, particularly focusing on the IHI. In line with the asymmetrical changes observed in information transfer following ccPAS stimulation, increasing interregional coupling in the root connecting the left M1 with the right M1 leads to an increase the inhibitory effect from the left M1 over the right M1 (i.e. reduced M1 cortical excitability), whereas the reversed stimulation (rM1-to-lM1 ccPAS) results in the opposite result pattern of reduced IHI – i.e. greater M1 excitability. In line with previous literature, the increased IHI following lM1-to-rM1 ccPAS did not appear immediately, but it built up with time emerging at about 20 minutes after the stimulation (Rizzo et al., 2009, 2011). Importantly, we observe a relationship between decreases in IHI following lM1-to-rM1 ccPAS and increases in the volume of information transferred between these two nodes following the stimulation. This relationship was circumscribed to the changes in information transfer in the high theta / low alpha oscillatory range (6-9 Hz). Previous neuromodulation studies report that the level of inhibition that the left M1 exerts over the right M1 varies with the degree of alpha rhythm phase synchronicity between these 2 regions, with the strongest IHI occurring when their phases are aligned (Stefanou et al., 2018). In the same vein, we found that manipulating the effective connectivity of the left M1-to-right M1 pathway increases volume information transfer in the 6–9 Hz range and its associated interregional synchronicity, which occurs in tandem with an increase in interhemispheric inhibition (IHI) from the left to the right M1. By directly linking changes in volume transfer to concurrent changes in corticospinal excitability and interhemispheric inhibition, we provide novel evidence for frequency-specific directional asymmetries underlying interhemispheric motor communication.

It is worth noting that the oscillatory changes in the beta, alpha, and theta bands are unlikely to result from volume conduction or changes in a distant site. This possibility was controlled for by using the weighted phase lag index, an unbiased estimator with low sensitivity to volume conduction and uncorrelated noise sources, ensuring that the changes in cortico-cortical coupling between the left and the right M1 were accurately measured. Importantly, with regards to cross-correlation, we only considered interhemispheric transmissions that take at least 8ms, thereby discarding any possible effects of volume conductance.

In summary, these findings illustrate a mechanistic link binding synaptic efficacy in short-range connections to their functional role as conveyors of information communication through oscillatory brain transmission. They highlight, for the first time, that interregional communication frequencies in the human lM1-to-rM1 pathway can be manipulated, leading to increases of information transfer instantiated by the phase alignment of the oscillatory architecture supporting motor control. The selective, frequency-specific changes in information volume and transfer speed observed after different ccPAS protocols may reflect the distinct functional roles of the two pathways interconnecting the left and right M1, particularly in a group of strongly right-handed participants. Taken together, these results are consistent with Hebbian-like spike-timing dependent long-term potentiation and depression (Koch et al., 2013); they also align common accounts of interregional communication, where precise temporal oscillatory phase alignment facilitates the efficient transfer of information across distant neural populations (Fries, 2005, 2015). This work paves the way for targeted neuromodulatory strategies aimed at enhancing motor function, with potential applications extending to other cognitive systems that exhibit similar patterns of asymmetrical connectivity. Future studies integrating behavioral measures will be critical to fully uncover the functional implications of these directional oscillatory changes, ultimately contributing to refined interventions for motor rehabilitation and performance optimization.

## Material and methods

### Participants

Sixty healthy, right-handed individuals participated across the three experiments: 20 participants in Experiment 1 (26.35 ± 5.21; 9; 81.25 ± 20.78), 20 participants in Experiment 2 (22.7 ± 4.18; 16; 80.31 ± 22.15) and 20 participants in Experiment 3 (25.6 ± 6.97; 16; 94.06 ± 9.61) (where numbers correspond to mean age ± SD; number of female participants, handiness mean ± SD; as measured by the Edinburgh handedness inventory (Veale, 2014). Participants were pseudo randomly allocated to each group. Due to technical difficulties during EEG data collection, 7 participants had to be excluding from the sample, which meant that the EEG data reported includes 17 participants in Experiment 1 (26.41 ± 5.58; 8; 83.09 ± 19.11), 18 participants in Experiment 2 (22.33 ± 3.73; 13; 79.51 ± 22.87), and 18 participants in Experiment 3 (25.94 ± 7.26; 14; 93.40 ± 9.93). All participants had no personal or familial history of neurological or psychiatric disease, were right-handed and gave written informed consent (Department of Psychology Research Ethics Committee, ETH1920-0135), were screened for adverse reactions to TMS and risk factors through a safety questionnaire (Rossi et al., 2021), and received either course credits or monetary compensation for their participation. Sample size was determined through power analyses for a repeated-measures, between-subject, design using medium effect sizes (f =.25) and >90% power to detect a significant effect (α level=.05). This computed sample size is in line with on previous studies that have used the same ccPAS protocol, along and in combination with EEG to manipulate connectivity between two human cortico-cortical regions in the motor control network (Fiori et al., 2018; Johnen et al., 2015; Sel et al., 2021).

### Experimental design

The three experiments started with a Baseline block, followed by a ccPAS period, and an Expression block (Figure 1). During Baseline and Expression blocks, TMS over rM1 and lM1 was delivered at rest. The effect exerted by the M1 conditioning pulse on the effect of the subsequent M1 test pulse was determined by contrasting motor-evoked potentials (MEPs) locked to the M1 test pulse recorded from the first dorsal interosseus (FDI) muscle, when either dual-site paired-pulse TMS was delivered over both right and left M1 (60 trials per block) or a single TMS pulse (60 trials per block) was applied to M1. Resting-state interactions between the right and the left M1 emerge at 8ms intervals (Rizzo et al., 2009, 2011). Precise interpulse timing is critical if both right and left M1 are to produce coincident influences on corticospinal activity. Therefore, the dual site paired pulses were applied with an interpulse interval (IPI) of 8 ms. In Experiment 1, the first pulse was always delivered to the lM1, followed by a pulse on the rM1 – i.e. lM1-to-rM1 stimulation. In Experiment 2 the pulse order was reversed, the rM1 pulse was followed by the lM1 pulse – i.e. rM1-to-lM1 stimulation. In Experiment 3, half of the participants received the first pulse on lM1, and half of the participants received the first pulse on rM1 (M1&M1 stimulation). Single pulses were delivered either on the rM1 or the lM1 in participants that underwent the lM1-to-rM1 ccPAS protocol or the rM1-to-lM1ccPAS protocol, respectively. For all participants and all groups, spTMS and ppTMS trials were pseudorandomized and counterbalanced within the same block. In addition, during Baseline and Expression blocks, we recorded resting EEG data while participants fixated on a white cross in the center of the computer screen for two 3-minute periods during eyes opened and during eyes closed – the order of these two periods was pseudorandomized and counterbalanced across participants. For the EEG data analysis, we focused on the EEG recorded during eyes opened.

In all three experiments, the ccPAS period that intervened between Baseline and Expression blocks consisted of 15 min of ccPAS over right and left M1 applied at 0.1 Hz (90 total stimulus pairings). In Experiment 1, the pulse applied to left M1 always preceded the pulse over right M1 (lM1-to-rM1 ccPAS), while the opposite was true in Experiments 2 (rM1-to-lM1ccPAS). In Experiments 1 and 2 the paired-pulses were delivered with an IPI of 8ms. In Experiment 3, the pulses were delivered almost simultaneously in a manner that does not follow Hebbian principles (1ms IPI - M1&M1 ccPAS), which served as an active control – half of the participants received the first pulse in the left M1 (followed by a pulse on the right M1 after 1ms), half of the participants received the first pulse in the right M1.

### TMS and electromyography recordings

TMS protocols were administered using two DuoMAG MP stimulators (BrainBox**®**), each connected to two 50 mm figure-of-eight coils. The M1 “scalp hotspot” for the right and the left M1 were defined using the resting motor threshold (rMT) method, and it was defined as the scalp location where TMS stimulation evoked 50μV amplitude in 5 out of 10 consecutive trials using the minimum intensity at rest. Scalp locations for right and left M1 were projected onto high-resolution standardized MRI image using frameless stereotactic neuronavigation (Brainsight; BrainBox**®**). Furthermore, scalp locations were used to project the trajectory into the cortex to define the “cortical hotspot” for all participants in Montreal Neurological Institute (MNI) coordinates (mean rM1 x = 49.03 ± 6.24, y = -8.48 ± 10.16, z = 63.54 ± 5.28 ; mean lM1, x = -47.23 ± 6.50, y = -10.81 ± 11.41, z = 63.56 ± 13.34; Figure 1) (Rizzo et al., 2011). These locations were similar to those reported previously (Buch et al., 2011; Davare et al., 2008; Johnen et al., 2015).

As in previous ccPAS studies (Rizzo et al., 2009, 2011), conditioning and test pulses of ccPAS protocol were set to 110% of rMT, eliciting single-pulse MEPs of ±1mV for each M1 (rM1: 61.7 ± 1.3; lM1: 59.24 ± 1.4; - numbers indicate percentage of mean and SE stimulator output) (Rossini et al., 1994). Given the variability of MEPs, TMS intensity was recalibrated during the initial 20 trials of the Expression block to ensure that the elicited MEPs remained within the ±1 mV range. TMS coils were held by the experimenters and were positioned tangential to the skull, with the M1 coil angled at ∼45° (handle pointing posteriorly),

Surface electromyography (EMG) was recorded from the right and left FDI with bipolar surface Ag-AgCl electrode montages. EMG signals were acquired at a sampling rate of 10000Hz and filtered (bandpass, 0.5 - 1000Hz), with additional hardwired 50Hz notch filtering (CED Humbug), and recorded using a CED D440-4 amplifier, a CED micro1401Mk.II A/D converter, and PC running Signal (7.07 version) (Cambridge Electronic Design). Motor evoked potentials (MEPs) were computed as the peak-to-peak amplitude difference. Trials in which MEPs were under 0.1mV (i.e. trials in which no MEP was elicited, mostly because of coil misplacement or missing stimulation due to other factors such as coil overheating) or over 6mV were discarded. Trials were excluded if the peak-to-peak FDI activity during the precontraction period, from 0.4 to 0.05 seconds before the pulse delivery, surpassed the 0.3 mV threshold (Experiment 1: 4.92 ± 6.29; Experiment 2: 6.40 ± 7.82; Experiment 3: 7.40 ± 7.53 - numbers indicate percentage of mean and SE of excluded trials for each Experiment). As the distribution of MEP amplitudes was positively skewed, they were log-transformed for further statistical analyses.

### MEP analysis

It is repeatedly shown that MEP amplitudes recorded at rest decrease when M1 TMS is preceded by a previous conditioning pulse over the contralateral M1 – i.e. the so-called interhemispheric inhibition (IHI) (Ferbert et al., 1992). In this line, we first computed the IHI for each participant as the ratio between the MEP amplitude elicited by the conditioned ppTMS and the mean MEP amplitude from the unconditioned spTMS. Higher IHI values correspond to a weaker inhibitory influence from the contralateral M1 (i.e., less inhibition), whereas lower values indicate a stronger inhibitory effect (Ferbert et al., 1992). Moreover, previous observations show that manipulating the pathway connecting the left and right M1 cortices impacts the IHI (Rizzo et al., 2009, 2011), and that these changes emerge sometime after the stimulation. Following previous evidence, we contrasted IHI values recorded at Baseline block *vs* those recorded at the Expression block using a repeated measures analysis of variance (ANOVA) with factors Experimental group (Experiment 1, Experiment 2, Experiment 3), Block (Expression, Baseline) and Time (first 20 trials, last 20 trials) – the factor time was implemented in the model to allow the investigation of whether the ccPAS influence on IHI evolved with time (Rizzo et al., 2009, 2011). To correct for non-sphericity in all repeated measure tests, Greenhouse-Geisser corrected results are reported. All post-hoc analyses were corrected for multiple comparisons using the Bonferroni Holms correction.

### EEG Recording and Analysis

EEG was recorded with sintered Ag/AgCl electrodes from 64 electrodes mounted equidistantly on an elastic electrode cap (64Ch-Standard-BrainCap for TMS; Brain Products**®**). Two electrodes measured the vertical (VEOG) and horizontal (HEOG) ocular activity, respectively. One electrode measured the electrocardiac (ECG) response. All electrodes were referenced to the AFz electrodes. Both eyes-closes and eyes-open resting state activity was recorded for 3 minutes at a rate of 1000Hz. All impedances were kept below 5kΩ, except for the first five participants, for whom the impedance levels were maintained below 10kΩ. Offline EEG analysis was performed using custom MATLAB scripts (version R2021a) and the EEGLAB toolbox (Delorme and Makeig, 2004), focusing on the EEG activity recorded during eyes-open resting state.

Resting-state data was filtered in the 0.5–100Hz band and a notch-filter at 50Hz was applied. Noisy channels were spherically interpolated, and bad segments data were eliminated. Next, an Independent Component Analysis (ICA) was implemented; ICA is an effective method widely employed for removal of EEG artifacts (Makeig et al., 2004). Components containing artefacts were removed from the data. Following this, we subtracted components related to vertical and horizontal eye movements (VEOG and HEOG) as well as cardiac activity (ECG). Then, the recording was re-referenced to the average of all electrodes, downsampled to 512Hz and a Laplacian transform was applied to the data using spherical splines. The Laplacian is a spatial filter that aids in topographic localization by attenuating artefacts attributable to volume conduction, rendering the data more suitable for performing connectivity analyses (Cohen, 2014). Finally, we extracted epochs lasting 2 seconds.

### Functional connectivity analysis

Functional connectivity between oscillatory activity of the two motor cortices was assessed by measuring their inter-areas phase relationship. Specifically, phase connectivity was estimated in the sensor space via the weighted phase-lag index (wPLI) (Vinck et al., 2011), a measure of the degree of synchronization between the signals. wPLI is based on the phase lag index (PLI) (Stam et al., 2007), which defines connectivity as the absolute value of the average sign of phase angle differences. In contrast to the PLI, wPLI gives maximal weighting to phase differences that are far from the real axis, and hence underweight all signals typically associated with artificial synchrony or volume conduction. wPLI values range between zero (i.e., random relationship between phases) and one (i.e., perfect phase synchronization). wPLI values were extracted, for each individual and for each block (Baseline and Expression), and for each individual frequency point (4-40Hz). The connectivity analysis of the EEG signal was focused on 24 electrodes distributed over frontocentral and centro-parietal areas, i.e. F1, F2, F3, F4, F5, F6, FC1, FC2, FC3, FC4, FC5, FC6, C1, C2, C3, C4, C5, C6, CP1, CP2, CP3, CP4, CP5, CP6, which cover the bilateral sensorimotor cortices (M1) and were organized into two distinct Regions of Interest (Left and Right ROI). We assessed the significance of spatial-frequency patterns in the extracted wPLI using a modified cluster-based permutation approach adapted from Groppe et al., 2011. In brief, this method first computes the test statistics, such as t-scores, for each time point and electrode site, comparing the experimental conditions. In our case, the t-scores refer to the difference in wPLI between Baseline and Expression Blocks. This method then identifies potential clusters of significant difference by setting an uncorrected alpha level (e.g., p < 0.05) and grouping adjacent time points and spatially neighboring sites exhibiting significant t-values that exceed this threshold. Clusters are defined by grouping the t-values that share the same sign (or by taking only one sign in a one-tailed test), and their cluster masses are then calculated by summing the t-values within each cluster to measure the strength and extent of the effect. In this way, in a two-tailed test, positive and negative clusters are obtained. At this point, it is necessary to generate the null distribution of the cluster masses. This involves shuffling the data—typically through 1,000 or more iterations—by randomly exchanging the wPLI measures recorded during the Baseline and Expression Blocks conditions. For each permutation, clusters are identified, and the largest cluster mass is recorded. These values form the null distribution, representing the cluster masses expected under the null hypothesis of no true effect. The final step is assessing cluster significance: any observed cluster mass exceeding the 95th percentile of the null distribution is deemed statistically reliable (i.e., beyond chance). This method is flexible and can be applied for both paired and independent sample testing, following the frameworks proposed by Bullmore et al. (1999) and Maris and Oostenveld (2007).

In our study, we devised a modified version of the Groppe’s approach (Groppe et al., 2011) because, unlike the original approach which is based on spatial adjacency of single electrodes, wPLI is a connectivity measure between two electrodes; therefore, the clustering had to be based on the adjacency of electrode pairs. To adapt the cluster-based method, which is typically designed for single electrodes, a definition of adjacency between electrode pairs was necessary. We adopted the most conservative rule: two pairs of electrodes were considered adjacent and thus belonging to the same cluster (same neuronal network), only if they shared at least one common electrode. For the analysis of the wPLI, we then compared the normalized data from the Baseline vs Expression blocks using the paired-sample version of the permutation method for each Experiment separately.

### Directionality analysis

Cross-correlation analysis is a widely used method for examining the presence and direction of causal relationships between signals, such as neural activity recorded from different brain regions. By measuring the correlation between the envelope of two signals as a function of time lag, this technique provides insight into their temporal relationships (Adhikari et al., 2010). However, as explained by El-Gohary and McNames (2007), cross-correlation can be misleading when signals exhibit strong autocorrelation, often resulting in broad correlation peaks centered at lag = 0, that obscure the true causal relationship. Therefore, to overcome this limitation, whitened cross-correlation is applied, utilizing whitening filters to remove autocorrelation effects and yield a clearer representation of connectivity (El-Gohary and McNames, 2007).

The results from the wPLI analysis allowed to identify 6 frequency windows where there was a significant increase of average functional connectivity (Experiment 1 – lM1-to-rM1 ccPAS: 6-9Hz, 13-16.5Hz, 19-25Hz; Experiment 2 – rM1-to-lM1 ccPAS: 3-5Hz, 6-11Hz, 14-19Hz, 24-30Hz).

To statistically assess differences between frequency bands for each experiment, a non-parametric analysis was performed using the Friedman test. When significant effects were observed, pairwise post hoc comparisons were conducted using Wilcoxon signed-rank tests. To control for multiple comparisons, p-values were adjusted using the False Discovery Rate (FDR) correction.

Within those frequency windows, we the cross-correlation between all the pairs of the ROIs and we performed again a cluster-based analysis to assess the directionality of the information flow. Following this data-driven approach, the brain signal was first filtered in the desired frequencies; then the envelope signal was evaluated and the whitening filter applied. The normalized cross-correlation was computed with the MATLAB function *xcorr*, here conventionally considering the right hemisphere EEG sensors as senders and the left hemisphere EEG sensors as receivers. Due to the bidirectional nature of the connection on the cortical region, the cross-correlation between the sender and the received signal is characterized by three main peaks (Figure 4A): a central peak (i.e. with 0s lag) representing the volume conductance component; a peak on the left side (i.e. with lag <0s) representing the information travelling from the sender to the receiver; a peak on the right side (i.e. with lag>0s) characterizing the information travelling back from the receiver to the sender. In communication between brain areas, the transmitter and receiver frequently switch roles (Hartung et al., 2016a, 2016b). To examine the directionality between two sensors capturing activity from two brain regions, the amplitude and lag of the side peaks were analyzed. Specifically, the amplitude is an index of volume of information transmitted, whereas the lag indexes the time it takes for information to travel from the transmitter to the receiver or vice versa, and therefore the lag is inversely proportional to the speed of the transmission (Adhikari et al., 2010). To assess the statistical significance of each peak amplitude and lag across the different electrode pairs, we applied the modified version of Groppe’s method as detailed above (2011). We used the paired sample version of the approach to test changes between the Baseline and the Expression blocks for each Experiment separately.

### EEG power spectrum analysis

In addition to examine functional synchronicity and cross-correlation, we measured changes in the amplitude (power) of the EEG signal. The power spectrum analysis of the EEG signal was focused on the electrodes where we analyzed changes in phase synchrony and cross-correlation (i.e. F1, F2, F3, F4, F5, F6, FC1, FC2, FC3, FC4, FC5, FC6, C1, C2, C3, C4, C5, C6, CP1, CP2, CP3, CP4, CP5, CP6). Using the MATLAB *pwelch* function (2 second Hanning window with 25% of overlap), we computed the power spectral density for each frequency bin between 1 and 40Hz. To investigate any effects of the ccPAS manipulation on power values, we used the Groppe and colleagues (2011) cluster-based permutation approach as defined earlier. Changes in the local spectral profile are unlikely as previous investigations have found no impact of manipulating interregional connections with ccPAS on the power of the EEG signal (e.g. Di Luzio et al., 2024; Sel et al., 2021).

### Evaluating the relationship between EEG cross-correlation and interhemispheric inhibition

Finally, we assessed the relationship between increases in volume and speed of communication and increases in the inhibitory influence of the left M1 over the right M1 following lM1-to-rM1 ccPAS in Experiment 1. To this aim, a Pearson’s correlation coefficient was computed between the changes in cross-correlation lag and volume measures (computed as the difference between Baseline block vs Expression block) and the changes in IHI obtained from contrasting the IHI measured in the Baseline vs the Expression block. For this analysis, we selected the electrode pairs who exhibited the greatest change in communication transfer following the ccPAS intervention. These were: C4-C5 for the frequency bands 6-9Hz and 13-16.5Hz, and the electrode pair C2-C3 for the 19-25Hz frequency window. FDR corrected for multiple comparison with alpha=0.05).

## Data availability statement

We have deposited all EEG and EMG raw data, as well as the code for the EEG data preprocessing implemented in MATLAB in an OSF repository. The link to the repository is: https://osf.io/w8ukb/overview

## Funding statement

Funding for this work was provided by the Academy of Medical Sciences Springboard Award (SBF008\1113), the Bial Foundation Grant (033/2024), and the Essex ESNEFT Psychological Research Unit for Behaviour, Health and Wellbeing (RCP15313) to Alejandra Sel, the University of Essex to Domiziana Falaschi (Chancellor’s PhD Scholarship) and to Cristina Del Prete (Faculty of Science and Health Partnership PhD Scholarship).

## Conflict of interest disclosure

The authors declare no conflict of interest

## Ethics approval statement

This study was approved by the Department of Psychology Research Ethics Committee – ethics number: ETH1920-0135

## Author contributions

A.S. designed the experiment; A.L., L.T., D.F. collected the data; A.L., L.T., C.D.P, V.D.F. analyzed the data; A.S., A.L., L.T., C.D.P., V.D.F., D.F. wrote the manuscript. All authors discussed the results and implications and commented on the manuscript at all stages.

## Acknowledgements

We thank Mr. Christian Rainer for his help in setting up the TMS equipment and the EMG recording system in the laboratory

## Supplementary Figures

**Figure S1:**
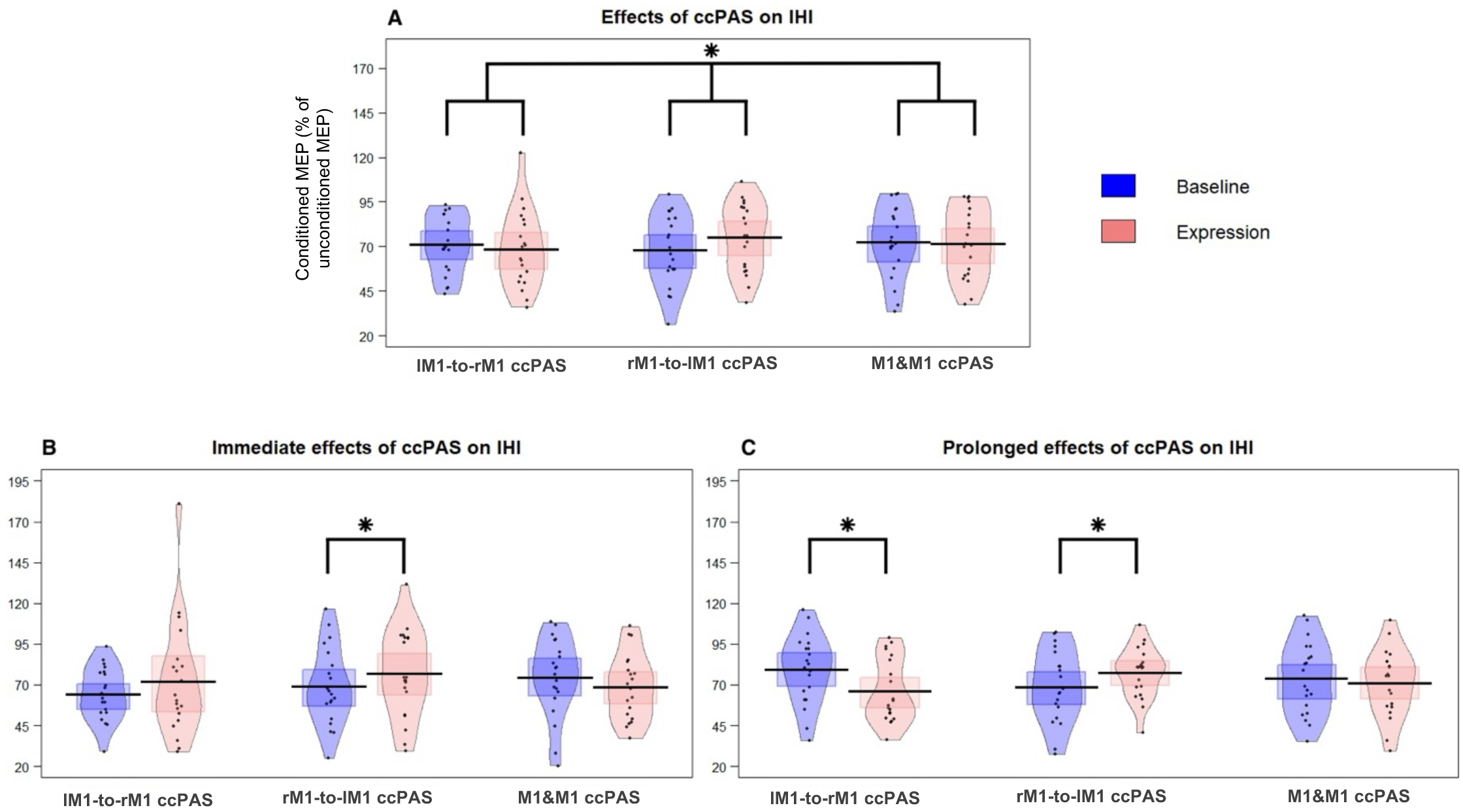
**A.** Overall effects of ccPAS on interhemispheric inhibition (IHI) across the three experiments – Group 1 (left, N=20), Group 2 (center, N=20), Group 3 (right=20). **B.** Immediate effects of ccPAS on IHI, predominantly observable in the rM1-to-lM1 ccPAS Group. **C**. Prolonged effects of ccPAS on IHI (observed in the last 20 trials), present in both Group 1 and Group 2, with opposite effects for each of these groups: in Group 1, we observe a reduced IHI after the ccPAS intervention, whereas in Group 2, the IHI enhance in the Expression block in comparison to the Baseline block. Blue bars represent the activity recorded at Baseline (i.e. before the ccPAS intervention); red bars indicate activity recorded after the ccPAS intervention. Shaded areas represent the 95% highest density interval; bars represent the mean value; the squared shared area represents the standard error of the mean; * denotes significant results with a p value <0.05

**Figure S2:**
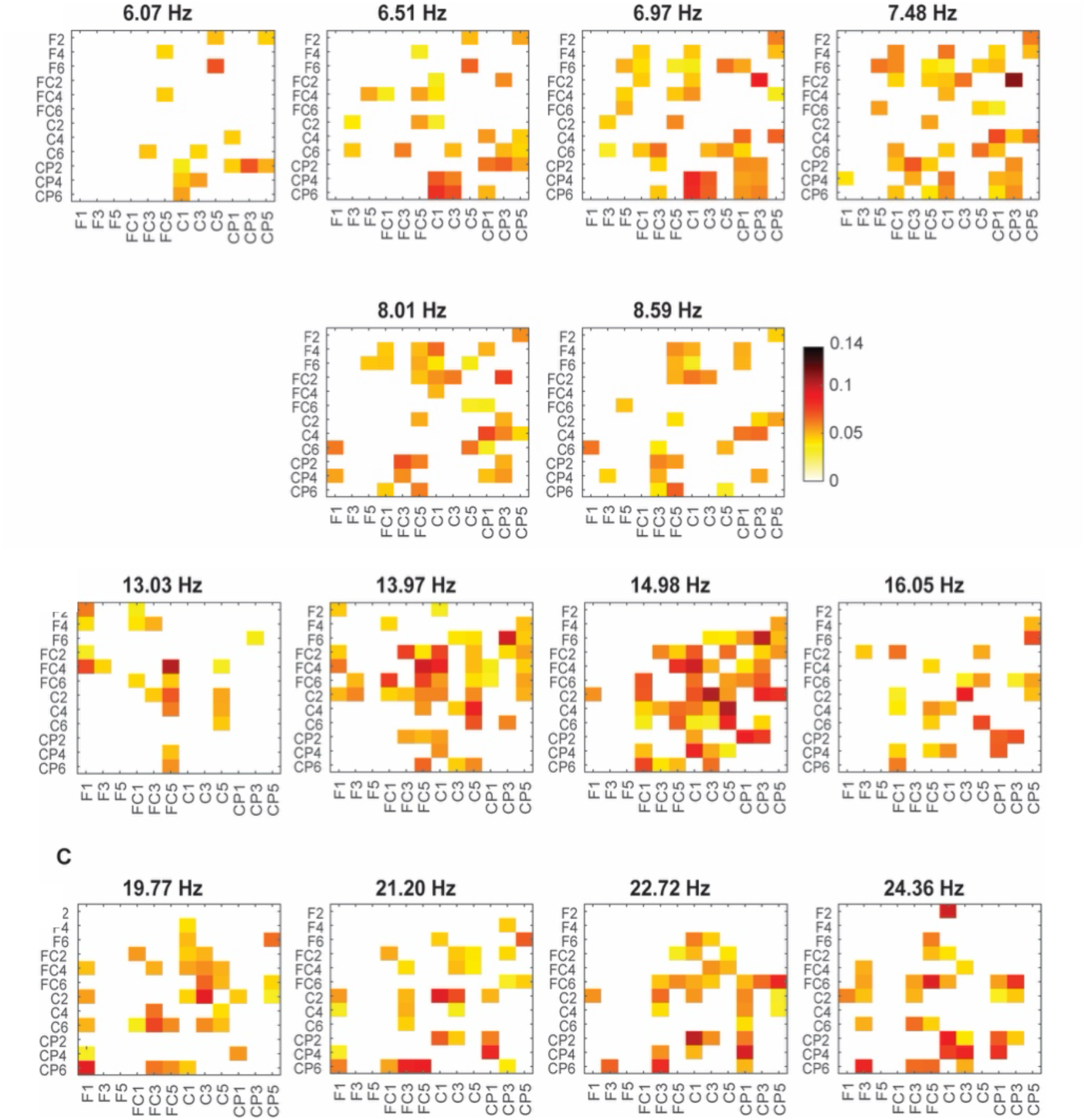
Effects of right M1-to-left M1 ccPAS over interregional connectivity Changes Before vs After ccPAS. 2-D representation of the changes in interregional coupling between the left M1 and the right M1 when contrasting activity recorded before and after lM1-to-rM1 ccPAS, averaged for each frequency range where significant wPLI changes were observed (N=17).

**Figure S3:**
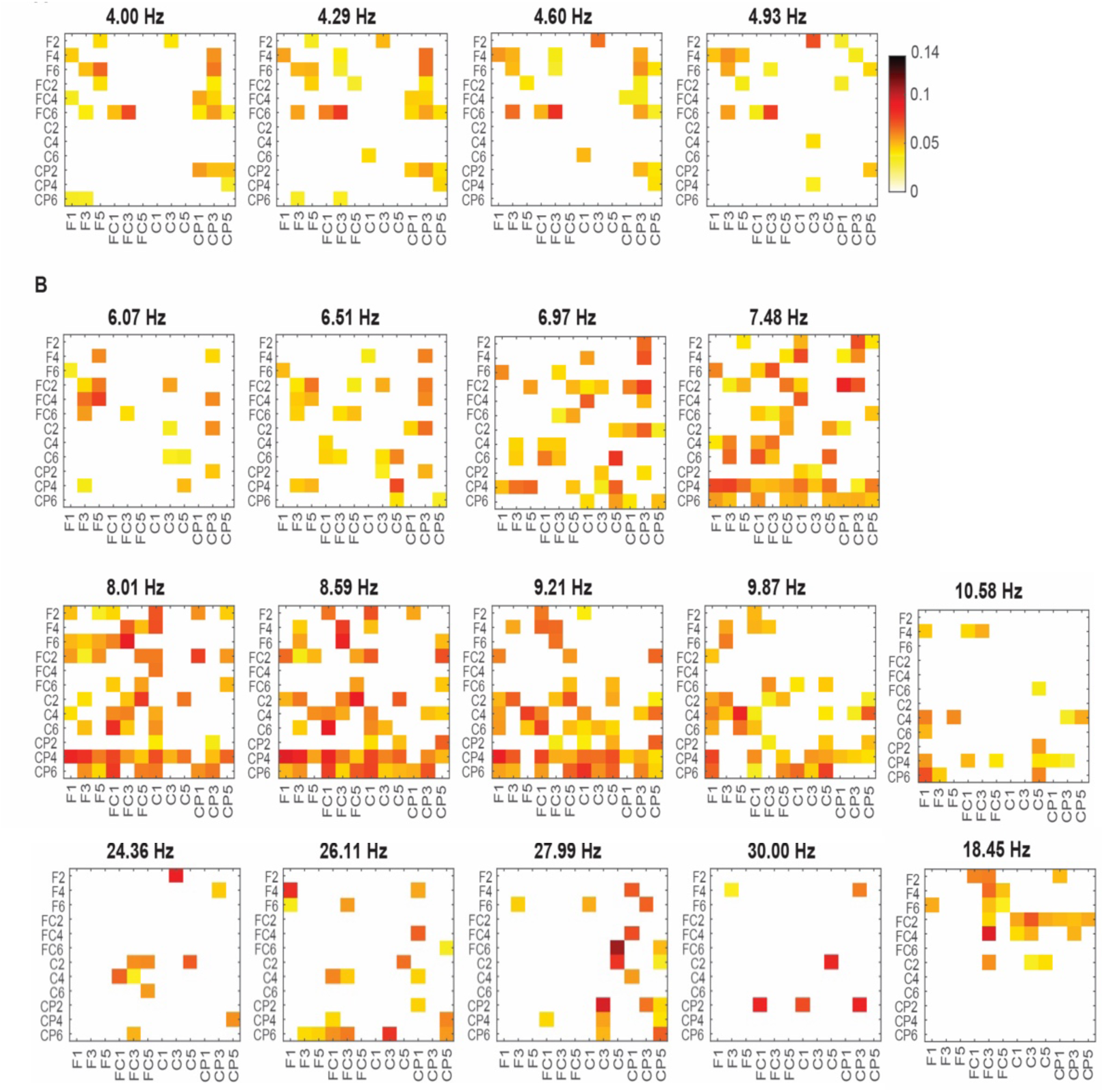
2-D representation of the changes in interregional coupling between the right M1 and the left M1 when contrasting activity recorded before and after rM1-to-lM1 ccPAS, averaged for each frequency range where significant wPLI changes were observed (N=18).

## Notes

https://osf.io/w8ukb/overview

### Competing Interest Statement

The authors have declared no competing interest.

